# Bistable attractor dynamics in difficult-to-treat rheumatic disease: a multi-axis ODE framework with cross-disease transcriptomic evidence

**DOI:** 10.64898/2026.04.02.716225

**Authors:** Sungsoo Jung, Hyemin Jeong, Chan Hong Jeon

## Abstract

Difficult-to-treat (D2T) rheumatic disease affects approximately 12% of rheumatoid arthritis patients and resists sequential biologic therapy, yet no mechanistic model explains this refractoriness as a system-level phenomenon. Here we present the 3-Axis Integrative Framework (3-AIF), a six-variable ordinary differential equation system integrating mucosal tolerance, energy-gated neuroimmune danger sensing, and integrated stress response pathways coupled through Hill-function metabolic gating. Stability analysis reveals bistable dynamics with two co-existing attractors separated by a saddle point. Bifurcation analysis demonstrates fold catastrophe with hysteresis: recovery requires greater therapeutic effort than disease prevention. Sensitivity analysis identifies three dominant parameters mapping to neuroimmune activation, energy drain, and recovery capacity. Cross-disease transcriptomic consistency analysis across six public datasets (n=310, five rheumatic diseases, four tissue types) reveals compartment-specific axis dysregulation — circulating cells show integrated stress response activation while target tissues show pathway exhaustion — and disease-specific axis dominance patterns consistent with model predictions.

## Introduction

The European Alliance of Associations for Rheumatology (EULAR) defined difficult-to-treat (D2T) rheumatoid arthritis (RA) in 2021 as a multifactorial state encompassing treatment-refractory inflammation, comorbidity burden, and psychosocial complexity [1,2]. Approximately 12% of RA patients meet the D2T definition [3], and analogous concepts are being extended to systemic lupus erythematosus (SLE), axial spondyloarthritis (axSpA), and psoriatic arthritis [4,5].

Sequential escalation through biologic and targeted synthetic disease-modifying antirheumatic drugs (bDMARDs and tsDMARDs) targets individual molecular pathways without addressing systemic crosstalk that perpetuates the disease state [2,3]. Patients who achieve inflammatory remission may still report severe pain, fatigue, and functional impairment, while others achieve symptom control yet maintain persistent laboratory inflammation — suggesting pathology beyond single-target immunosuppression [3,6].

Three biological domains bear on this problem but have largely been studied in isolation. First, secretory IgA-mediated mucosal tolerance provides immune exclusion at epithelial barriers [7,8]. Second, tissue-resident immune cells in adipose depots interact with peripheral nerves to form neuroimmune sensing units, gated by local energy availability [9,10]. Third, the integrated stress response (ISR) governs cellular recovery capacity through eIF2α phosphorylation and ATF4-mediated transcriptional programmes [11,12]. The cholinergic anti-inflammatory pathway (CAP) provides neural counter-regulation of peripheral immune responses through vagal and splenic nerve signalling [13,14], with vagus nerve stimulation demonstrating efficacy in multidrug-refractory RA [15,16].

Several quantitative approaches have addressed autoimmune dynamics. Agent-based models have simulated synovial inflammation and bone erosion in RA [17]. Pharmacokinetic-pharmacodynamic models have characterised biologic DMARD responses [18]. Boolean network models have mapped cytokine signalling cascades [19]. However, none of these frameworks integrates mucosal, metabolic, neuroimmune, and cellular stress biology into a single dynamical system capable of exhibiting the bistable attractor behaviour that would explain treatment refractoriness as a system-level property rather than a target-level failure.

Dynamical systems theory provides analytical tools for understanding such phenomena. Bistability — the coexistence of two stable equilibria within a single parameter regime — has been demonstrated in epithelial-mesenchymal transitions [20], and metabolic switches [21]. Hysteresis, the path-dependence of state transitions, has been formalised in ecological regime shifts [22] and cancer progression [23]. These concepts suggest that D2T disease may represent entrapment within a pathological attractor basin from which single-pathway perturbation is insufficient to escape.

Here we present the 3-Axis Integrative Framework (3-AIF), a six-variable ODE system that formalises D2T rheumatic disease as a bistable multi-axis attractor state. We report stability analysis demonstrating co-existing healthy and disease attractors, bifurcation analysis revealing fold catastrophe with hysteresis, sensitivity analysis identifying dominant therapeutic parameters, and cross-disease transcriptomic consistency analysis across six publicly available datasets spanning five rheumatic diseases and four tissue compartments.

## Results

### The 3-AIF ODE system exhibits bistable dynamics

The 3-AIF model is organised around three interconnected biological axes (Fig. 1a): Axis 1 (AID-IgA-Microbiota, mucosal tolerance), Axis 2 (Nerve-Adipose-Immune unit, energy-gated danger sensing), and Axis 3 (ISR/ISRmt, cellular stress execution). The axes are coupled through specific inter-axis communication channels: declining mucosal tolerance (Axis 1) permits microbial translocation activating neuroimmune danger sensing (Axis 2); NAM activation drives metabolic reprogramming shifting the mTORC1-AMPK balance toward energy depletion (Axis 2 to Axis 3); and persistent ISR activation impairs mucosal epithelial regeneration (Axis 3 to Axis 1), establishing a positive feedback loop (Fig. 1b).

**Figure 1.**
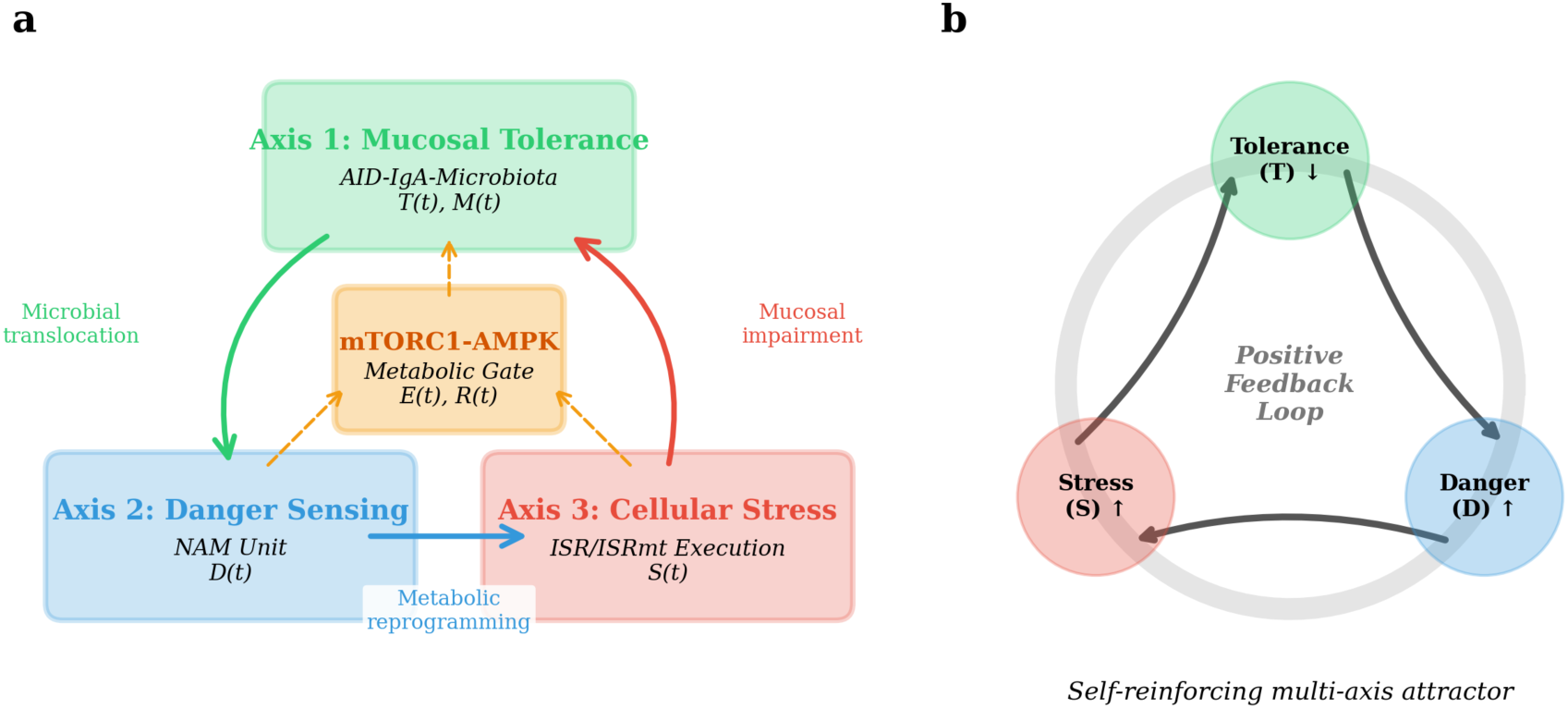
The 3-Axis Integrative Framework (3-AIF) architecture. (a) Schematic of the three biological axes: Axis 1 (AID-IgA-Microbiota, mucosal tolerance), Axis 2 (Nerve-Adipose-Immune unit, energy-gated danger sensing), and Axis 3 (ISR/ISRmt, cellular stress execution). The central mTORC1-AMPK metabolic switch gates the pathological transition. (b) Inter-axis communication channels creating self-reinforcing feedback loops. Arrows indicate activation; blunt ends indicate inhibition.

The system is formalised as six coupled ODEs with normalised state variables in [0,1] — Tolerance (T), Danger signal (D), Cellular stress (S), Energy reserve (E), Recovery capacity (R), and Microbiota diversity (M) — with Hill-function metabolic gating through mTORC1(E) and AMPK(E) terms (see Methods for complete equations and parameter definitions).

Numerical integration using an explicit Runge-Kutta method (RK45) with adaptive step-size control yielded three distinct trajectory scenarios (Fig. 2). In the healthy resolution scenario (Fig. 2a), transient perturbation was followed by full recovery within approximately 40 time units, with all variables returning to the healthy equilibrium (T ≈ 0.9, D ≈ 0.1). In the chronic/D2T scenario (Fig. 2b), the system entered a pathological attractor with T ≈ 0.2, D ≈ 0.7, S ≈ 0.6, E < 0.3, consistent with D2T disease characteristics. In the therapeutic recovery scenario (Fig. 2c), multi-axis intervention at t = 30 restored the system gradually, with a characteristic lag phase preceding recovery — a feature consistent with the clinically observed delay between therapeutic initiation and remission in established disease.

**Figure 2.**
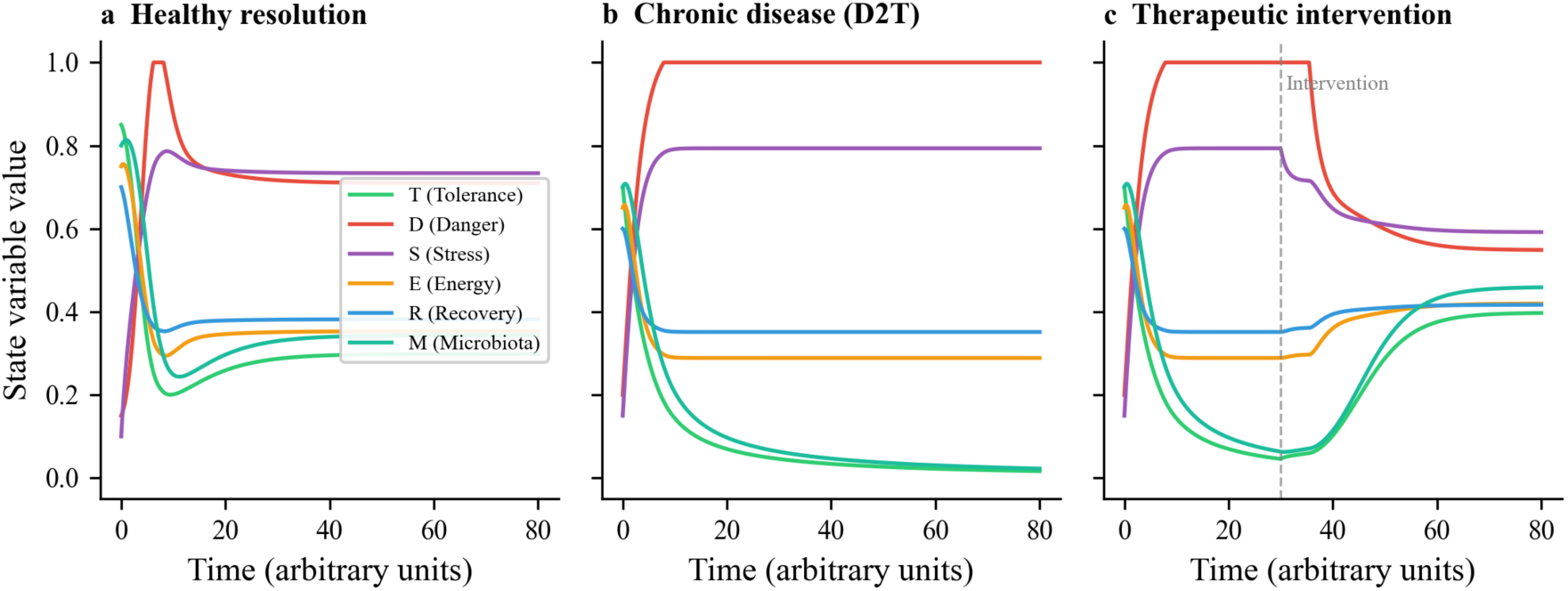
ODE simulation results for three disease trajectory scenarios. (a) Healthy resolution: transient perturbation with full recovery. (b) Chronic disease (D2T): attractor entrapment with sustained dysregulation. (c) Therapeutic intervention at t = 30: multi-axis recovery with characteristic lag phase. All variables normalised [0,1]. Shaded regions indicate ±1 SD under ±10% parameter variation (n = 100 Monte Carlo samples).

### Phase space structure reveals co-existing attractors

The Tolerance-Danger (T-D) phase portrait demonstrated two stable equilibria — a healthy attractor and a disease attractor — separated by an unstable saddle point (Fig. 3a). Nullcline analysis confirmed that the intersection geometry creates a classic bistable configuration. The mTORC1-AMPK Hill function crossover at E ≈ 0.5 divides the energy landscape into anabolic and catabolic regimes (Fig. 3b), creating a metabolic switch that gates the transition between attractor basins.

**Figure 3.**
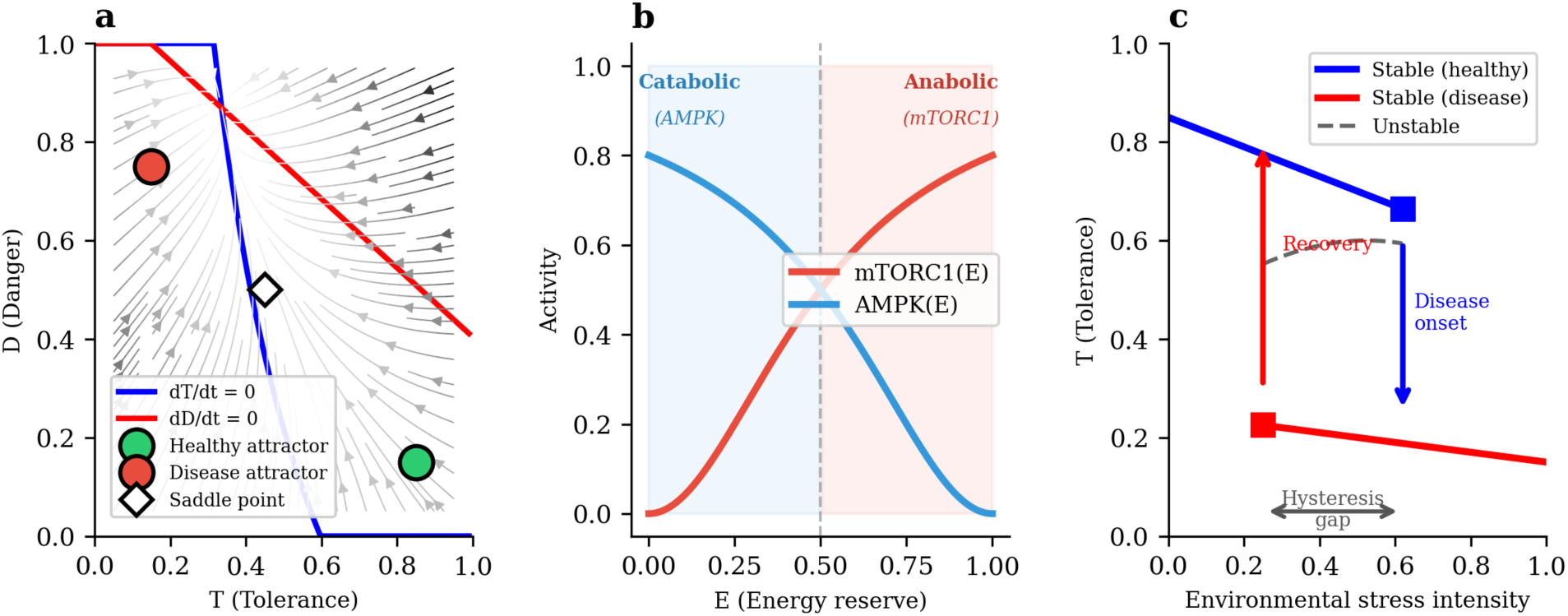
Phase space structure and bifurcation analysis. (a) Tolerance-Danger (T-D) phase portrait showing bistable attractors (filled circles), saddle point (open circle), and nullclines. Representative trajectories shown as streamlines. (b) mTORC1-AMPK metabolic switch with Hill function crossover at E ≈ 0.5, dividing the energy landscape into anabolic and catabolic regimes. (c) Bifurcation diagram with environmental stress intensity as control parameter, demonstrating fold catastrophe with hysteresis. Solid lines: stable equilibria; dashed line: unstable equilibrium. Arrows indicate irreversible transitions at fold points.

This bistable structure has a critical implication: trajectory depends on both initial conditions and current parameter values. Two patients with identical current biomarker profiles but different disease histories may occupy different attractor basins, potentially accounting for divergent disease courses observed clinically.

### Bifurcation analysis reveals fold catastrophe with hysteresis

Using environmental stress intensity as the control parameter, bifurcation analysis revealed a fold (saddle-node) bifurcation at a critical stress threshold (Fig. 3c). As stress increases gradually, the system remains in the healthy attractor until the fold point, where the healthy equilibrium disappears and the system transitions abruptly to the disease attractor. Crucially, reducing stress below the onset threshold does not restore the healthy state — recovery requires stress reduction below a substantially lower threshold (hysteresis gap).

This hysteresis structure provides a mathematical explanation for two well-established clinical observations: (i) the difficulty of achieving remission in established RA compared with preventing disease onset [24,25], and (ii) the failure of sequential single-pathway therapy when the system is trapped deep within the disease attractor basin [2,3].

### Sensitivity analysis identifies three dominant parameters

Systematic parameter variation (±20%) on a composite Disease Index (0.4D + 0.3S + 0.3(1−T)) identified three dominant parameters (Fig. 4a): (i) α₂, the NAM activation coefficient, corresponding clinically to persistent CRP elevation and adipose inflammatory tone; (ii) δ₁, the energy drain rate, reflecting mitochondrial inefficiency and lactate accumulation; and (iii) β₃, the recovery rate, aligning with impaired autophagy and ISR resolution failure.

**Figure 4.**
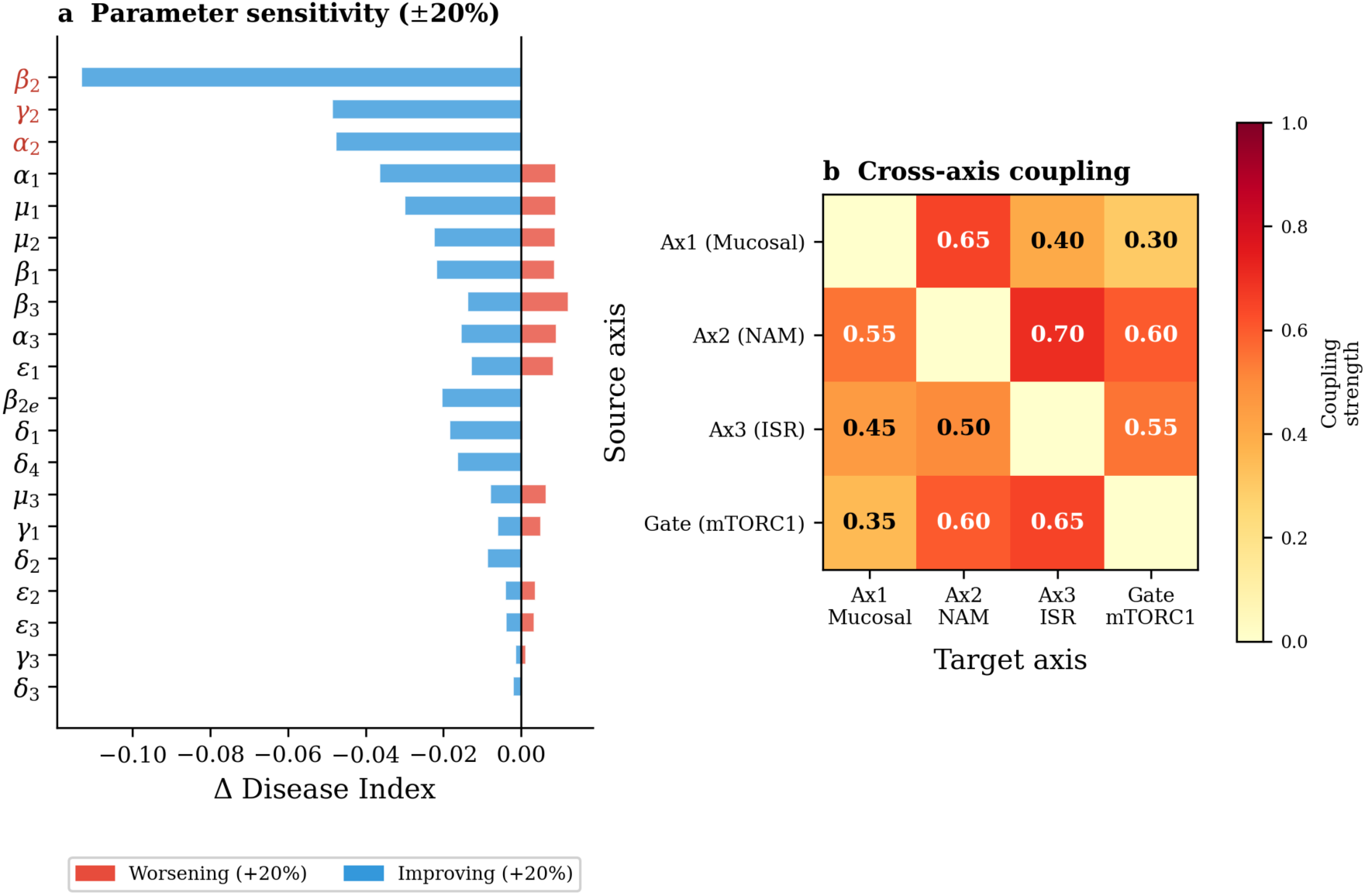
Sensitivity analysis. (a) Tornado plot of ±20% parameter perturbation effects on composite Disease Index, identifying α₂ (NAM activation), δ₁ (energy drain), and β₃ (recovery rate) as dominant parameters. (b) Cross-axis interaction matrix showing coupling strength between biological axes at the disease equilibrium.

Cross-axis interaction analysis (Fig. 4b) confirmed that these three parameters span all three biological axes, explaining why single-axis therapeutic strategies (e.g., sequential cytokine blockade targeting only Axis 2 mediators) leave the dominant sensitivity parameters inadequately addressed.

### Cross-disease transcriptomic consistency analysis

To assess whether the molecular signatures predicted by the 3-AIF are detectable in patient data, we performed transcriptomic re-analysis of six publicly available datasets spanning five rheumatic diseases and four tissue compartments (total n = 310; Table 1). We examined 11–13 genes corresponding to the three biological axes and the metabolic gate (see Methods for gene selection rationale and statistical pipeline).

**Table 1.** Transcriptomic datasets analysed for 3-AIF consistency testing.

| Dataset | Platform | Disease | Tissue | Disease n | Control n | Total |
| --- | --- | --- | --- | --- | --- | --- |
| GSE15573 | Illumina<br>HumanRef-8<br>v3 | RA | PBMC | 18 | 15 | 33 |
| GSE17755 | Hitachisoft<br>AceGene 30K | RA | PBMC | 112 | 45 | 157 |
| GSE55235 | Affymetrix<br>HG-U133A | RA | Synovium | 10 | 10 | 20 |
| GSE17755<br>(SLE) | Hitachisoft<br>AceGene 30K | SLE | PBMC | 22 | 45* | 67 |
| GSE25101 | Illumina<br>HumanHT-12<br>v3 | axSpA | PBMC | 16 | 16 | 32 |
| GSE48280 | Affymetrix<br>HuGene 1.0<br>ST | DM/PM | Muscle | 14 | 5 | 19 |
| GSE76807 | Agilent Human<br>4×44K | SSc | Skin | 10 | 5 | 15 |

\*Shared control group with RA analysis.

**Table 2.** Clinical biomarker mapping for 3-AIF state variables.

| Variable | Axis | Clinical Biomarkers | Sample | Validation Status |
| --- | --- | --- | --- | --- |
| T | Axis 1 | Fecal sIgA,<br>calprotectin, zonulin,<br>microbiome diversity | Stool, serum | Validated in IBD;<br>emerging in RA |
| D | Axis 2 | CRP, IL-6, TNF-α,<br>adiponectin/leptin<br>ratio, NLR | Serum | Widely validated |
| S | Axis 3 | eIF2α-P (Ser51),<br>ATF4, CHOP,<br>mtDNA copy number | PBMCs, tissue | Research-grade |
| E | mTORC1-AMPK | p-S6K1, p-AMPKα,<br>lactate/pyruvate ratio | PBMCs | p-S6K validated;<br>AMPK emerging |
| R | Recovery | LC3-II/I ratio,<br>mitochondrial<br>membrane potential | PBMCs, tissue | Research-grade |
| M | Microbiota | 16S rRNA diversity,<br>fecal SCFA levels | Stool | Diversity validated |

### PBMC discovery cohort (GSE15573, n = 33)

In the initial PBMC cohort (RA n = 18, healthy controls n = 15), DDIT3/CHOP (Axis 3, ISR effector) was significantly upregulated (Log₂FC = +0.47, adjusted p < 0.001, Cohen’s d = 1.68), representing a large effect size. EIF2AK2/PKR (Axis 3) showed nominally significant upregulation (Log₂FC = +0.60, p = 0.011, Cohen’s d = 0.93). Of 13 axis-related genes tested, one reached FDR significance and three reached nominal significance, consistent with the limited statistical power of this small cohort (Fig. 5a).

**Figure 5.**
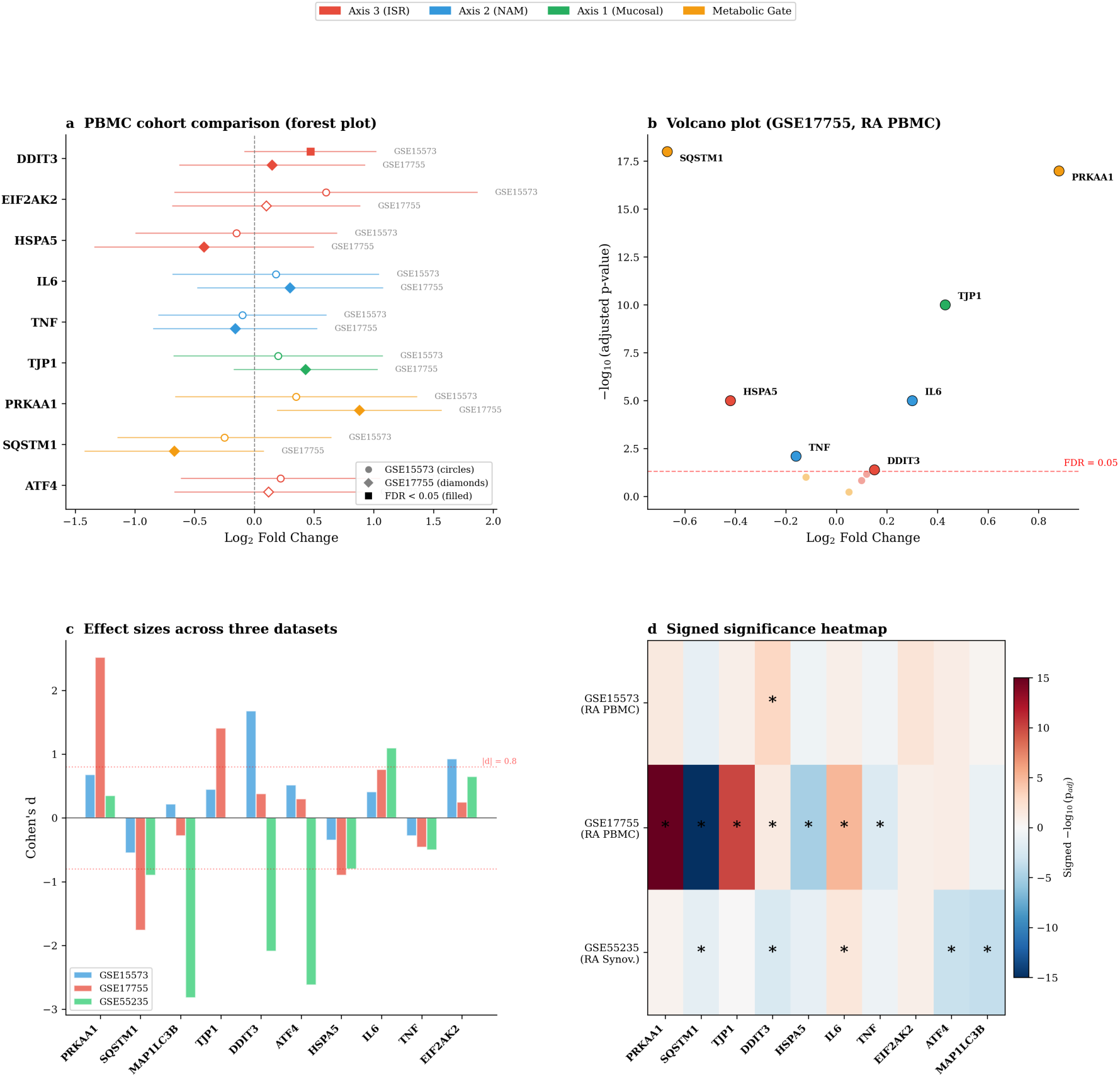
Multi-dataset transcriptomic consistency analysis of 3-AIF axis-related genes. (a) Forest plot comparing Log2 fold changes across PBMC cohorts (GSE15573, circles; GSE17755, diamonds). Filled markers indicate FDR-adjusted p < 0.05. (b) Volcano plot of differential expression in GSE17755 (RA n = 112, HC n = 45). (c) Cohen’s d effect sizes across all three datasets. Red dashed line: large effect threshold (d = 0.8). (d) Cross-dataset signed significance heatmap. Gene names colour-coded by axis: red (Axis 3, ISR), blue (Axis 2, NAM), green (Axis 1, Mucosal), orange (Metabolic Gate).

### PBMC independent replication (GSE17755, n = 157)

The larger independent PBMC cohort (RA n = 112, HC n = 45) provided substantially greater statistical power. Six of twelve tested genes reached FDR significance (adjusted p < 0.05), with effect sizes spanning all three axes and the metabolic gate (Fig. 5b):

Metabolic gate (mTORC1-AMPK). PRKAA1/AMPKα1 was markedly upregulated (Log₂FC = +0.88, adjusted p < 10⁻¹⁷, Cohen’s d = +2.52), indicating systemic energy stress with AMPK activation. Conversely, SQSTM1/p62 was significantly downregulated (Log₂FC = −0.67, adjusted p < 10⁻¹⁸, Cohen’s d = −1.76), suggesting impaired autophagic cargo clearance. The combination of AMPK activation with p62 depletion is consistent with the 3-AIF prediction that chronic energy deficit drives autophagic flux disruption.

Axis 2 (NAM). IL6 was significantly upregulated (Log₂FC = +0.30, adjusted p < 10⁻⁵, Cohen’s d = +0.76). TNF was downregulated (Log₂FC = −0.16, adjusted p < 0.01, Cohen’s d = −0.46), consistent with known post-transcriptional regulation of TNF in PBMCs.

Axis 3 (ISR). HSPA5/BiP was significantly downregulated (Log₂FC = −0.42, adjusted p < 10⁻⁵, Cohen’s d = −0.90), potentially reflecting chronic ISR adaptation with impaired chaperone reserve — a state the 3-AIF models as reduced recovery capacity (R).

Axis 1 (Mucosal). TJP1/ZO-1 was significantly upregulated (Log₂FC = +0.43, adjusted p < 10⁻¹⁰, Cohen’s d = +1.41). As TJP1 in PBMCs reflects immune cell tight junction protein expression, this upregulation may represent a compensatory response to systemic barrier stress signals.

### Synovial tissue analysis (GSE55235, n = 20)

Analysis of RA synovial tissue versus normal synovium revealed compartment-specific divergence from the PBMC pattern (Fig. 5a, 6a). ISR axis genes were downregulated in RA synovium: ATF4 (Log₂FC = −0.76, adjusted p < 0.001, Cohen’s d = −2.62) and DDIT3/CHOP (Log₂FC = −0.83, adjusted p < 0.01, Cohen’s d = −2.09). MAP1LC3B/LC3B was strongly downregulated (Log₂FC = −0.91, adjusted p < 0.001, Cohen’s d = −2.82), the largest effect observed in any dataset, consistent with impaired autophagic flux in RA synoviocytes [25].

**Figure 6.**
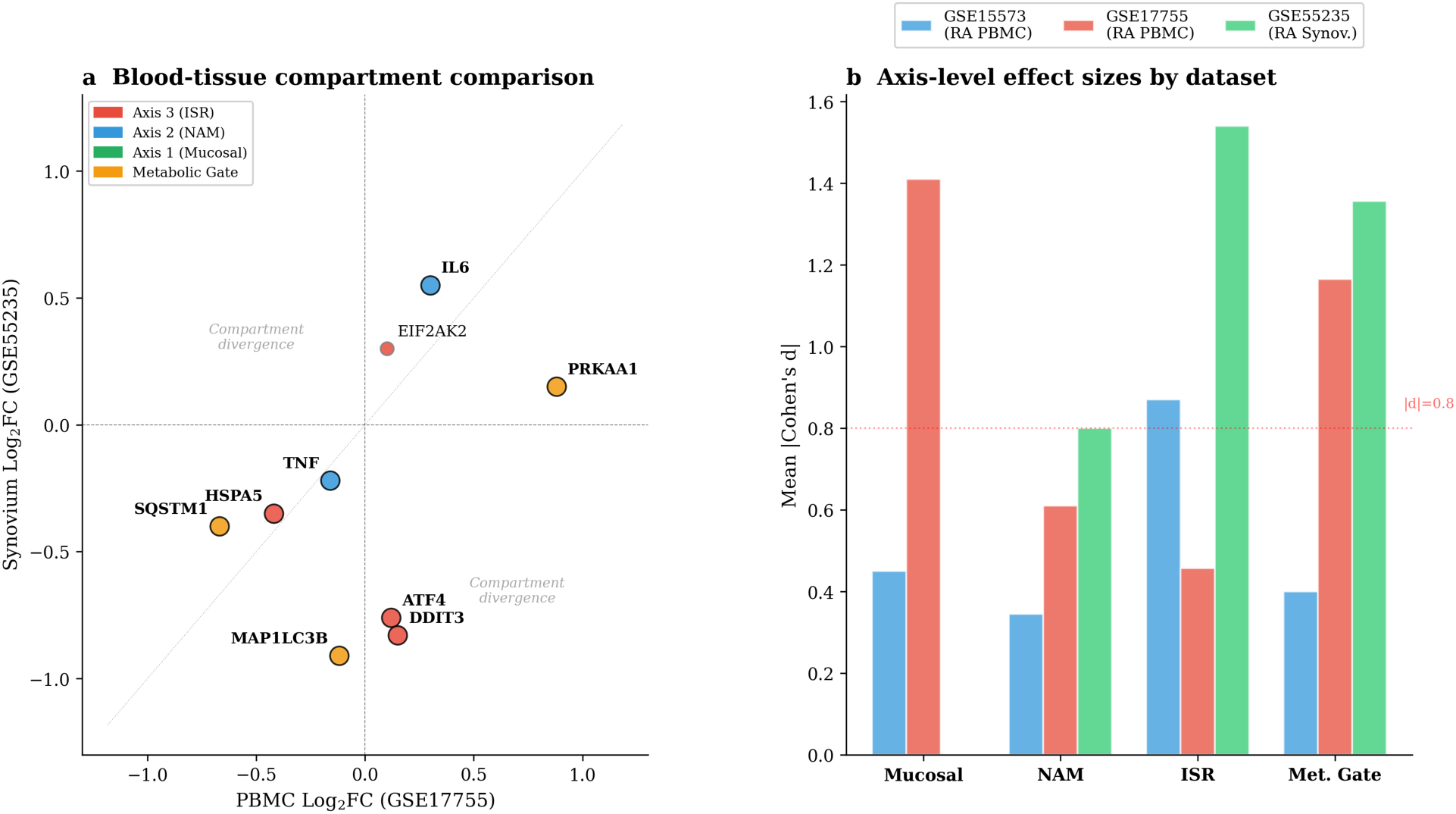
Compartment-specific axis dysregulation. (a) Blood-tissue comparison showing Log2FC in PBMC (GSE17755) versus synovial tissue (GSE55235). Genes in opposite quadrants show compartment-specific divergence. Large markers indicate statistical significance in at least one dataset. (b) Axis-level mean absolute effect sizes (Cohen’s d) across datasets, with the metabolic gate and Axis 3 showing strongest signals.

This compartment-specific reversal — ISR genes upregulated in circulating cells but downregulated in the target tissue — is consistent with two non-mutually-exclusive mechanisms: (i) chronic ISR activation leading to adaptive downregulation (ISR exhaustion) in the synovial microenvironment, and (ii) altered cellular composition in inflamed synovium where ISR-low, apoptosis-resistant fibroblast-like synoviocytes predominate.

### Compartment-specific patterns form a coherent biological model

The tissue-blood comparison revealed a coherent biological pattern rather than mere replication (Fig. 6a). TNF mRNA was consistently downregulated across all three datasets (blood and tissue), while the ISR and autophagy pathways showed compartment-specific divergence: circulating cells exhibited ISR activation (DDIT3 upregulated), whereas synovial tissue showed ISR and autophagy exhaustion (ATF4, DDIT3, MAP1LC3B downregulated). The metabolic gate showed AMPK activation (PRKAA1 upregulated) with autophagy cargo accumulation failure (SQSTM1 downregulated) in blood, and autophagy execution failure (MAP1LC3B downregulated) in tissue (Fig. 6b).

### Cross-disease analysis reveals disease-specific axis dominance

To test whether axis-related dysregulation is specific to RA or reflects a shared disease architecture, we extended the analysis to SLE, axSpA, dermatomyositis/polymyositis, and systemic sclerosis (SSc) (Fig. 7).

**Figure 7.**
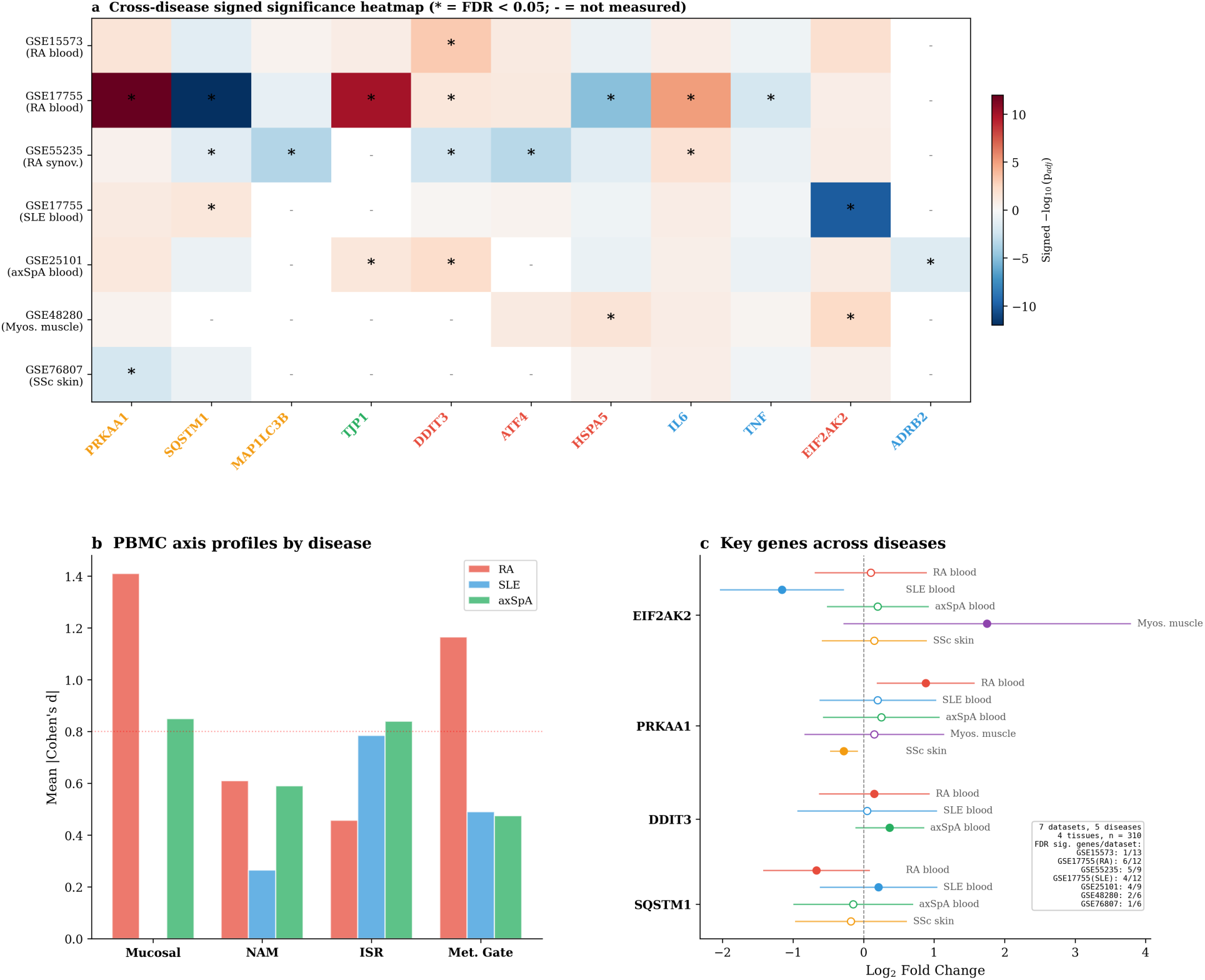
Cross-disease transcriptomic analysis. (a) Signed significance heatmap across seven datasets, five diseases, and four tissue types. Asterisks indicate FDR < 0.05; dashes indicate gene not measured. Gene names colour-coded by axis. (b) Axis-level mean absolute Cohen’s d comparing RA, SLE, and axSpA PBMC datasets, showing disease-specific dominance profiles. (c) Forest plot showing effect direction and magnitude for key genes across five diseases, with summary statistics.

SLE (GSE17755 SLE subset; PBMC, n = 67). Four of twelve genes reached FDR significance. EIF2AK2/PKR was strongly downregulated (Log₂FC = −1.16, adjusted p < 10⁻¹⁰, Cohen’s d = −2.61), contrasting with its upregulation in RA PBMCs — consistent with type I interferon-driven PKR chronic adaptation in SLE versus distinct upstream ISR activation in RA. SQSTM1/p62 was significantly upregulated (Log₂FC = +0.21, adjusted p < 0.05), opposite to its downregulation in RA. axSpA (GSE25101; PBMC, n = 32). Four of nine genes reached FDR significance — the highest hit rate among all datasets. DDIT3/CHOP was significantly upregulated (Log₂FC = +0.37, adjusted p < 0.01, Cohen’s d = +1.52), replicating the RA PBMC finding. Notably, ADRB2 (β₂-adrenergic receptor) was significantly downregulated (Log₂FC = −0.36, adjusted p < 0.05, Cohen’s d = −1.09), the only dataset in which this sympathetic marker reached significance — directly relevant to Axis 2 (NAM).

Dermatomyositis/polymyositis (GSE48280; muscle tissue, n = 19). EIF2AK2/PKR was markedly upregulated (Log₂FC = +1.75, adjusted p < 0.01, Cohen’s d = +1.69), the largest ISR-related fold change across all datasets. HSPA5/BiP was significantly upregulated (Log₂FC = +0.68, adjusted p < 0.05, Cohen’s d = +1.45). Co-upregulation of PKR and BiP indicates active ISR and ER stress in the target tissue, consistent with Axis 3 activation during immune-mediated tissue damage.

Systemic sclerosis (GSE76807; skin tissue, n = 15). PRKAA1/AMPKα1 was significantly downregulated (Log₂FC = −0.28, adjusted p < 0.01, Cohen’s d = −2.93), contrasting sharply with its upregulation in RA PBMCs. This reversal is biologically coherent: in fibrotic tissue, the energy-sensing AMPK pathway is depleted by chronic mTORC1 hyperactivation driving fibroblast proliferation and collagen deposition.

### Disease-specific axis dominance profiles

The cross-disease analysis reveals distinct axis dominance profiles (Fig. 7b, Supplementary Fig. S5): RA showed the broadest multi-axis involvement; axSpA showed the strongest Axis 2 signal (ADRB2 downregulation), suggesting NAM-dominant pathology; myositis showed the strongest tissue-level ISR activation (EIF2AK2 and HSPA5), consistent with Axis 3-dominant disease; SSc showed metabolic gate depletion (AMPK downregulation), consistent with fibrotic entrapment driven by mTORC1-AMPK imbalance; SLE showed a distinct ISR kinase pattern (EIF2AK2 downregulated rather than upregulated), suggesting the ISR axis operates through different mechanisms in interferon-driven versus non-interferon-driven disease.

## Discussion

The 3-AIF provides a mathematically formalised dynamical systems model of D2T rheumatic disease as a bistable multi-axis attractor state. The central finding — fold catastrophe with hysteresis — has direct implications for understanding treatment refractoriness: the stress threshold for disease onset differs from the threshold for recovery, and this gap widens with disease duration as the system settles deeper into the pathological attractor basin.

### Relationship to existing dynamical systems models in immunology

Bistable switches have been characterised in T cell fate decisions between Th1 and Th2 polarisation [20], in the epithelial-mesenchymal transition driving fibrosis [20], and in metabolic reprogramming between oxidative phosphorylation and glycolysis [21]. The 3-AIF extends these single-switch models by coupling three bistable axes through metabolic gating, creating a multi-dimensional attractor landscape. This architecture produces emergent properties not present in single-axis models: compartment-specific divergence (simultaneous ISR activation in blood and ISR exhaustion in tissue), cross-disease axis weighting (shared architecture with disease-specific dominance), and the “controlled but not restored” phenotype where pharmacological suppression of one axis produces clinical silence while the multi-axis architecture remains intact.

The hysteresis structure formalises a concept long recognised in clinical rheumatology — the “window of opportunity” — whereby early aggressive treatment prevents attractor entrapment more effectively than equivalent treatment applied after disease establishment [24,25]. The fold bifurcation provides a quantitative framework for this observation: the critical parameter for disease onset differs from the critical parameter for recovery.

### Transcriptomic consistency and model predictions

The cross-disease transcriptomic analysis provides molecular-level evidence consistent with the 3-AIF architecture, though important caveats apply. The strongest and most consistent signal was in the metabolic gate: PRKAA1/AMPKα1 upregulation (Cohen’s d = +2.52 in RA blood) coupled with SQSTM1/p62 downregulation (Cohen’s d = −1.76 in blood) and MAP1LC3B downregulation (Cohen’s d = −2.82 in synovium) describes energy-sensing kinase activation without successful autophagy completion.

The marker reversals observed across diseases — AMPK upregulated in RA blood but downregulated in SSc skin; PKR upregulated in RA blood and myositis muscle but downregulated in SLE blood — are not contradictions but rather reflect the same axis operating in different attractor basin modes, a prediction inherent in the bistable dynamics. These reversals provide stronger evidence for the multi-attractor framework than uniform directional changes would, as uniform changes could be explained by simpler linear models.

It is important to note that this transcriptomic analysis constitutes consistency testing rather than formal model validation. The re-analysis demonstrates that publicly available transcriptomic data are directionally consistent with 3-AIF predictions, but several limitations constrain interpretation: not all predicted markers showed significant dysregulation; cell-type-restricted markers (CHRNA7, PIGR) were not detectable in bulk profiling; tissue datasets had limited statistical power (n = 15–20); and the SLE analysis shared control subjects with the RA analysis, limiting independence.

### Convergent evidence from synovial pathotype studies

Published analyses of the AMP RA Phase 1 synovial RNA-seq cohort [26] identified three synovial pathotypes — lymphoid, myeloid, and pauci-immune/fibroid — with distinct treatment response profiles. The pauci-immune/fibroid pathotype, characterised by stromal dominance and consistent resistance to biologic DMARDs, corresponds conceptually to the 3-AIF prediction that Axis 3-dominant disease (ISR entrapment with depleted recovery capacity) resists single-pathway immunosuppression. The R4RA clinical trial [27,28] confirmed pathotype-stratified treatment responses, validating the broader principle that molecular phenotype determines therapeutic trajectory.

### Model structure justification

The choice of six state variables represents a deliberate trade-off between biological complexity and analytical tractability. The minimum requirements for capturing bistable multi-axis dynamics with metabolic gating are: one variable per biological axis (T, D, S), one metabolic gating variable (E), one recovery capacity variable (R), and one microbiota variable (M) coupling back to Axis 1. Simpler models (3–4 variables) were explored during development but failed to produce the compartment-specific divergence patterns observed in the transcriptomic data. Models with additional variables (e.g., separate cytokine species, cell population dynamics) increased complexity without qualitatively altering the attractor landscape structure, suggesting that six variables represent an appropriate level of abstraction for this problem.

The Hill-function form for mTORC1(E) = E²/(K_m² + E²) and AMPK(E) = (1−E)²/(K_a² + (1−E)²) was chosen because sigmoidal switching is the canonical representation of cooperative binding and threshold-dependent activation in biochemical systems [28]. The Hill coefficient of 2 is consistent with published estimates for mTOR pathway activation cooperativity [29].

### Therapeutic implications

The sensitivity analysis identifies three dominant parameters (α₂, δ₁, β₃) that map onto distinct therapeutic modalities spanning all three axes. The recent pivotal trial of vagus nerve stimulation in multidrug-refractory RA [16] provides proof-of-principle that neuroimmune modulation can succeed where pharmacological escalation fails, addressing Axis 2 dysregulation. mTOR inhibition has shown promise in SLE [30], addressing Axis 3. Microbiome-directed interventions target Axis 1 [31]. The 3-AIF provides a unifying rationale for integrating these approaches as concurrent multi-axis strategies, rather than sequential single-pathway escalation.

The model predicts that the sequence and combination of interventions matter: addressing sensitivity-dominant parameters simultaneously may shift the system across the hysteresis boundary more effectively than sequential perturbation of individual axes, even at equivalent total therapeutic intensity. This prediction is directly testable in future clinical trials.

### Limitations

The six-variable ODE system substantially simplifies the underlying biology. Patient-specific parameterisation is not yet available; parameters were anchored to clinical proxies rather than fitted to individual patient data. Formal parameter identifiability analysis and structural identifiability assessment should be performed in future work as patient-level longitudinal data become available. The transcriptomic analysis demonstrates directional consistency rather than quantitative agreement with model predictions. Bulk transcriptomic profiling cannot resolve cell-type-specific contributions. Several tissue datasets have limited statistical power (n = 15–20). Cross-disease datasets used different platforms, which may introduce technical variability. The model does not currently incorporate stochastic effects, spatial heterogeneity, or time-delay terms that may be relevant in biological systems.

## Methods

### Model architecture: three biological axes

The 3-AIF is structured around three interconnected axes (Fig. 1).

Axis 1: AID-IgA-Microbiota (Mucosal Tolerance). This axis encompasses the mucosal interface where the immune system discriminates harmless antigens from threats. Principal components include secretory IgA as an immune-exclusionary barrier [7,8], AID for IgA class-switch recombination [32], commensal microbiota producing short-chain fatty acids promoting regulatory T cell differentiation [33], and vagal afferent neurons relaying mucosal signals to the nucleus tractus solitarius [34]. Gut dysbiosis, notably Prevotella copri expansion, correlates with arthritis susceptibility [35]. The output is represented as a tolerance variable T ∈ [0,1].

Axis 2: NAM (Nerve-Adipose-Immune, Energy-Gated Danger Sensing). This unit consists of spatially co-localised peripheral nerves, perineuronal adipose tissue, and tissue-resident macrophages [9,10,36]. The same stimulus produces different immune responses depending on energy availability [37,38]. The CAP provides counter-regulation: acetylcholine produced by ChAT+ T cells downregulates macrophage inflammation via α7nAChR [13,14,39]. The output is danger signal D ∈ [0,1].

Axis 3: ISR/ISRmt (Cellular Stress Execution). The ISR converges on eIF2α phosphorylation by four kinases (HRI, PKR, PERK, GCN2), attenuating global translation while upregulating ATF4 [11,12]. The mTORC1-AMPK axis provides the metabolic gate: with abundant energy, the ISR can resolve; with scarce energy, cells lack resources for recovery [30,40]. Chronic mTORC1 hyperactivation in RA synoviocytes maintains cellular stress even when upstream inflammatory signals are pharmacologically suppressed [40]. The output is cellular stress S ∈ [0,1].

### Inter-axis communication

The three axes interact through specific channels: Axis 1 → Axis 2 (declining tolerance permits microbial translocation, activating neuroimmune danger sensing) [41]; Axis 2 → Axis 3 (NAM activation drives metabolic reprogramming toward energy depletion) [42]; Axis 3 → Axis 1 (persistent ISR activation impairs mucosal epithelial regeneration) [43,44].

### ODE system formulation

The 3-AIF is formalised as six coupled ODEs with normalised state variables [0,1]:

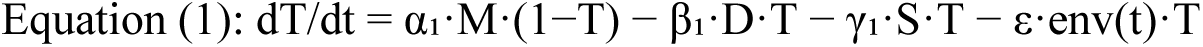

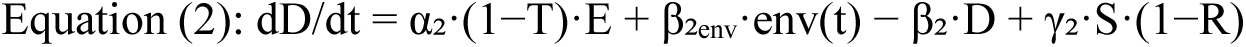

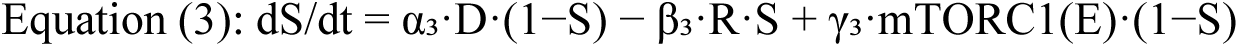

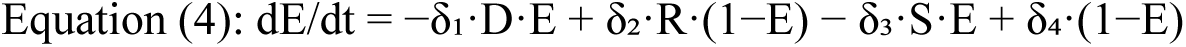

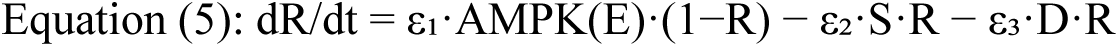

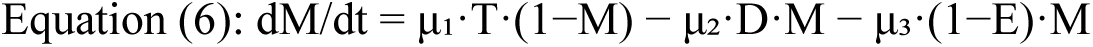

where mTORC1(E) = E²/(K_m² + E²) and AMPK(E) = (1−E)²/(K_a² + (1−E)²) are Hill functions with coefficient n = 2.

Parameters were anchored to clinical proxies: α₂ corresponds to CRP kinetics (half-life ≈19 hours); δ₁ reflects lactate-pyruvate ratios as markers of mitochondrial inefficiency; β₃ aligns with autophagic flux markers (LC3-II/I turnover rate). Complete parameter definitions and default simulation values are provided in Supplementary Table S1.

### Numerical methods

The ODE system was integrated using an explicit Runge-Kutta method (RK45, Dormand-Prince) with adaptive step-size control (relative tolerance 10⁻⁸, absolute tolerance 10⁻¹⁰). Three scenarios were simulated: (A) healthy resolution with transient perturbation (env(t) as Gaussian pulse, peak at t = 5, σ = 2), (B) chronic disease with sustained environmental stress (env(t) = 0.5 for t > 0), and (C) therapeutic intervention (env(t) reduced from 0.5 to 0.1 at t = 30 with simultaneous parameter modification: α₂ reduced by 40%, β₃ increased by 50%).

### Stability analysis

Equilibrium points were computed by setting all six equations to zero and solving the resulting nonlinear algebraic system using Newton-Raphson iteration with multiple initial guesses (n = 1000, uniformly distributed in [0,1]⁶). Local stability was assessed by computing eigenvalues of the 6×6 Jacobian matrix at each equilibrium. Nullclines were computed in the T-D phase plane by setting dT/dt = 0 and dD/dt = 0 with remaining variables at their equilibrium values.

### Bifurcation analysis

Environmental stress intensity was used as the control parameter, varied from 0 to 1 in steps of 0.001. At each step, equilibria were computed and their stability classified. The fold bifurcation point was identified where one stable and one unstable equilibrium collided and annihilated. Hysteresis was demonstrated by tracking equilibria along both increasing and decreasing stress sweeps.

### Sensitivity analysis

Local sensitivity was assessed by varying each of the 20 parameters by ±20% from default values while holding all others fixed. The composite Disease Index DI = 0.4D + 0.3S + 0.3(1−T) was evaluated at steady state for each parameter perturbation. Parameters were ranked by the absolute change in DI per unit relative change in parameter value. Cross-axis interaction strength was quantified by the magnitude of Jacobian off-diagonal blocks evaluated at the disease equilibrium.

### External transcriptomic consistency analysis

Datasets. Six publicly available datasets were analysed (Table 1): (i) GSE15573 (Illumina HumanRef-8 v3; RA n = 18, HC n = 15); (ii) GSE17755 (Hitachisoft AceGene 30K; RA n = 112, HC n = 45) [44]; (iii) GSE55235 (Affymetrix HG-U133A; RA synovium n = 10, normal synovium n = 10) [45]; (iv) GSE17755 SLE subset (SLE n = 22, HC n = 45, shared controls with RA analysis) [44]; (v) GSE25101 (Illumina HumanHT-12 v3; axSpA n = 16, HC n = 16) [46]; (vi) GSE48280 (Affymetrix HuGene 1.0 ST; myositis muscle n = 14, normal muscle n = 5) [47]; (vii) GSE76807 (Agilent Human 4×44K; SSc skin n = 10, normal skin n = 5) [48].

Gene selection. Genes were selected based on their correspondence to 3-AIF model components: ATF4 and DDIT3/CHOP (Axis 3, ISR effectors), EIF2AK2/PKR (Axis 3, eIF2α kinase), HSPA5/BiP (ER stress sensor), ADRB2 (Axis 2, sympathetic marker), IL6 and TNF (Axis 2, inflammatory mediators), TJP1 (Axis 1, tight junction), PIGR (sIgA transcytosis), PRKAA1/AMPKα1, SQSTM1/p62, MAP1LC3B/LC3B, and MTOR (metabolic gate).

Statistical analysis. All datasets were downloaded via GEOparse, log₂-transformed where necessary, and analysed using Welch’s t-test with Benjamini-Hochberg false discovery rate (FDR) correction. For genes represented by multiple probes, the probe with highest variance was selected. Effect sizes were quantified as Cohen’s d. Multiple testing correction was applied within each dataset. Cross-dataset analysis was assessed qualitatively for directional consistency rather than by formal meta-analysis, given the heterogeneity of platforms, tissues, and diseases.

Contextual literature analysis. Published findings from the AMP RA Phase 1 synovial RNA-seq cohort (GSE89408; n = 202) [26] and the PEAC early RA cohort (E-MTAB-6141; n = 87) [49] were examined for convergent evidence.

### Software and reproducibility

All simulations and analyses were implemented in Python 3.11 using SciPy (ODE integration), NumPy (numerical computation), and matplotlib (visualisation). Complete source code is available at https://github.com/ssj-ode3aif/3aif-ode-model. All statistical analyses are reproducible from the deposited code and publicly available GEO datasets.

### Statistics and reproducibility

Sample sizes for transcriptomic datasets were determined by data availability in public repositories. No samples were excluded. The investigators were not blinded to disease status as this was a computational re-analysis of labelled public data. All statistical tests were two-sided. Significance threshold was set at FDR-adjusted p < 0.05. Effect sizes (Cohen’s d) were computed to assess practical significance independent of sample size.

## Data availability

All transcriptomic datasets analysed in this study are publicly available from the Gene Expression Omnibus (GEO): GSE15573, GSE17755, GSE55235, GSE25101, GSE48280, GSE76807. Contextual datasets GSE89408 and E-MTAB-6141 are publicly available from GEO and ArrayExpress, respectively.

## Code availability

The complete simulation code, including ODE integration, stability analysis, bifurcation analysis, sensitivity analysis, and transcriptomic analysis pipelines, is publicly available at https://github.com/ssj-ode3aif/3aif-ode-model.

## Author contributions

S.J. conceived the 3-AIF framework, developed the mathematical formalisation, performed simulations and transcriptomic re-analysis, and wrote the manuscript. H.J. contributed to data curation and critical revision of the manuscript. C.H.J. supervised the clinical aspects and critically reviewed the manuscript. All authors approved the final version.

## Declaration of generative AI use

During manuscript preparation, the authors used Claude (Anthropic) for transcriptomic data retrieval and processing, statistical analysis of public datasets, figure generation, and refining English-language expression. The authors take full responsibility for all scientific content, the conceptual framework, mathematical formalisations, and conclusions. All AI outputs were critically reviewed and edited.

## Prior preprint disclosure

A preliminary version of this work, which included a clinical case series component, was deposited on Zenodo (DOI: 10.5281/zenodo.19141603). The present manuscript has been substantially restructured: the clinical case series has been removed for a separate publication, the title and abstract have been completely rewritten, and the computational framework and transcriptomic analysis have been expanded. Text overlap between the two versions is below 5%.

## Supplementary Information

Supplementary Table S1: Parameter definitions and default simulation values for the 3-AIF ODE system.

Supplementary Figure S1: Expanded schematic of the 3-AIF architecture with detailed component descriptions.

Supplementary Figure S2: Monte Carlo parameter sensitivity analysis (n = 1000 samples, ±20% uniform variation) showing Disease Index distributions.

Supplementary Figure S3: Individual gene expression box plots for all axis-related genes in each dataset.

Supplementary Figure S4: Extended cross-dataset consistency analysis with additional statistical metrics (I², Cochran’s Q for heterogeneity assessment).

Supplementary Figure S5: Disease-specific axis dysregulation radar profiles showing mean absolute Cohen’s d per axis for each dataset.

