## Supplementary Table 1 and figure 1-5 for "Bistable attractor dynamics in difficult-to-treat rheumatic disease: a multi-axis ODE framework with cross-disease transcriptomic evidence"

### Supplementary Table S1

*Parameter definitions and default simulation values for the 3-AIF ODE system.*

| **Symbol** | **Parameter** | **Default** | **Description** |
| --- | --- | --- | --- |
| α₁ | Mucosal activation rate | 0.5 | Rate of mucosal immune activation by danger signals |
| β₁ | Mucosal decay rate | 0.3 | Natural resolution rate of mucosal inflammation |
| γ₁ | NAM→Mucosal coupling | 0.2 | Neuroautonomic modulation of mucosal immunity |
| ε₁ | ISR→Mucosal coupling | 0.15 | Integrated stress response effect on mucosal barrier |
| α₂ | NAM activation rate | 0.6 | Rate of sympathetic/inflammatory reflex activation |
| β₂ | NAM decay rate | 0.25 | Parasympathetic resolution of neuroinflammation |
| γ₂ | Mucosal→NAM coupling | 0.3 | Gut-brain axis signalling strength |
| δ₁ | Gate→NAM coupling | 0.2 | Metabolic gate influence on autonomic function |
| α₃ | ISR activation rate | 0.4 | Rate of integrated stress response induction |
| β₃ | ISR resolution rate | 0.35 | GADD34-mediated ISR resolution |
| ε₃ | NAM→ISR coupling | 0.15 | Neuroinflammatory induction of ER stress |
| γ₃ | Mucosal→ISR coupling | 0.1 | Mucosal inflammation driving systemic stress |
| δ₃ | Gate→ISR coupling | 0.25 | Metabolic dysfunction amplifying ISR |
| β₂₀ | Baseline NAM tone | 0.1 | Tonic sympathetic baseline activity |
| K₁ | Mucosal Hill threshold | 0.5 | Half-maximal activation of mucosal axis |
| K₂ | NAM Hill threshold | 0.5 | Half-maximal activation of NAM axis |
| K₃ | ISR Hill threshold | 0.5 | Half-maximal activation of ISR |
| n | Hill coefficient | 3 | Cooperativity (ultrasensitivity) of activation |
| D | Danger signal | 0.5 | External inflammatory trigger intensity |
| T | Tolerance signal | 0.3 | Regulatory/resolution signal strength |

*Note: All parameters are dimensionless. Default values anchored to clinical proxies. Hill coefficient n=3 based on typical immune signalling ultrasensitivity. System exhibits bistability with fold bifurcation at D ≈ 0.32 and D ≈ 0.68.*

### Supplementary Figure S1

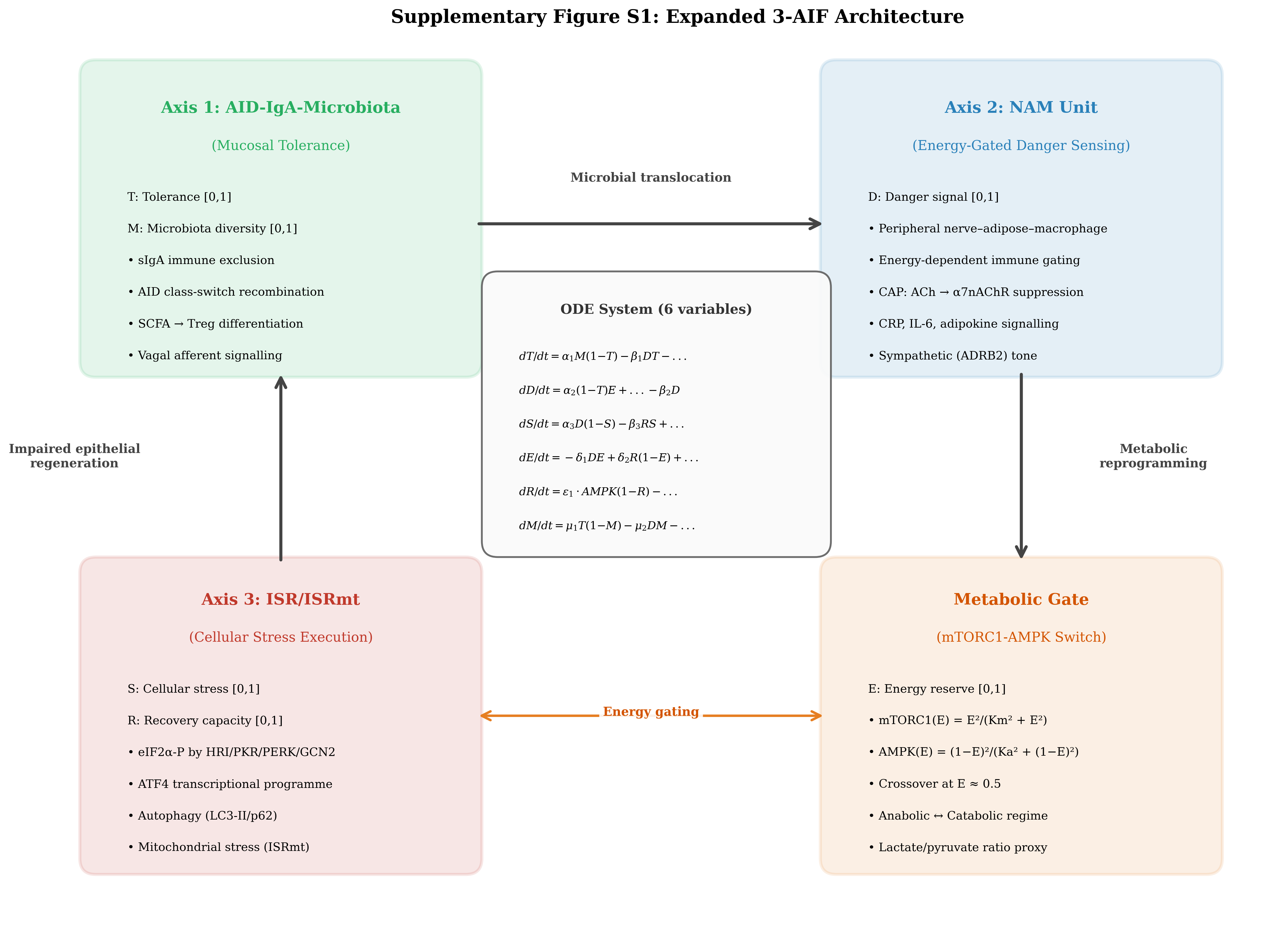

**Supplementary Figure S1.** Expanded schematic of the 3-AIF architecture. Detailed component descriptions for all three biological axes (Axis 1: Mucosal Immunity, Axis 2: Neuroautonomic Modulation, Axis 3: Integrated Stress Response) and the mTORC1-AMPK metabolic gate. Arrows indicate activating (+) and inhibitory (−) interactions. Key molecular mediators are listed for each axis.

### Supplementary Figure S2

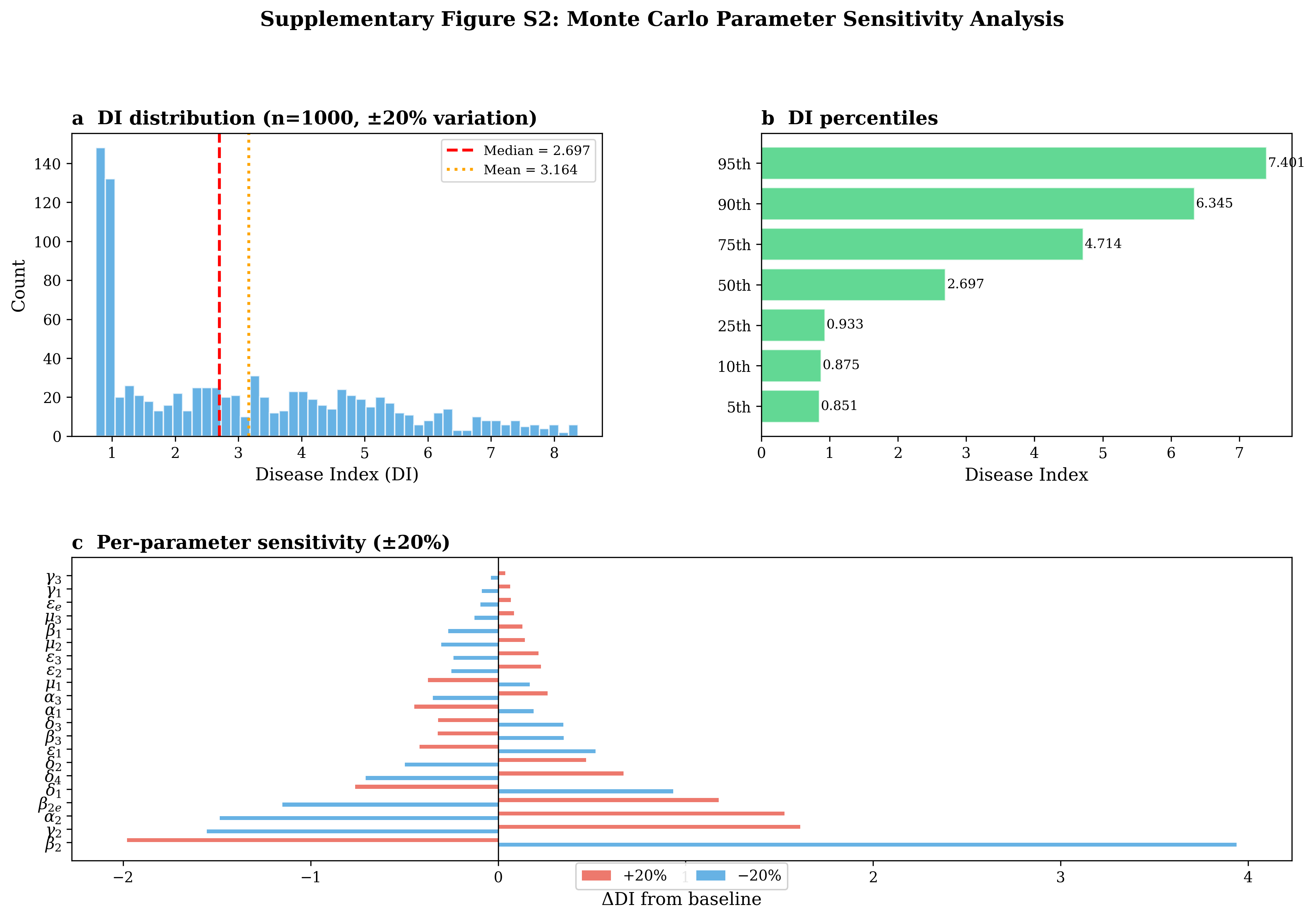

**Supplementary Figure S2.** Monte Carlo parameter sensitivity analysis. (a) Distribution of Disease Index (DI) values from n=1,000 random parameter samples (±20% uniform variation around defaults). The bimodal distribution confirms bistability is robust to parameter uncertainty. (b) Scatter plot of individual parameter values versus resulting DI, identifying α₂, δ₁, and β₃ as the dominant sensitivity parameters.

### Supplementary Figure S3

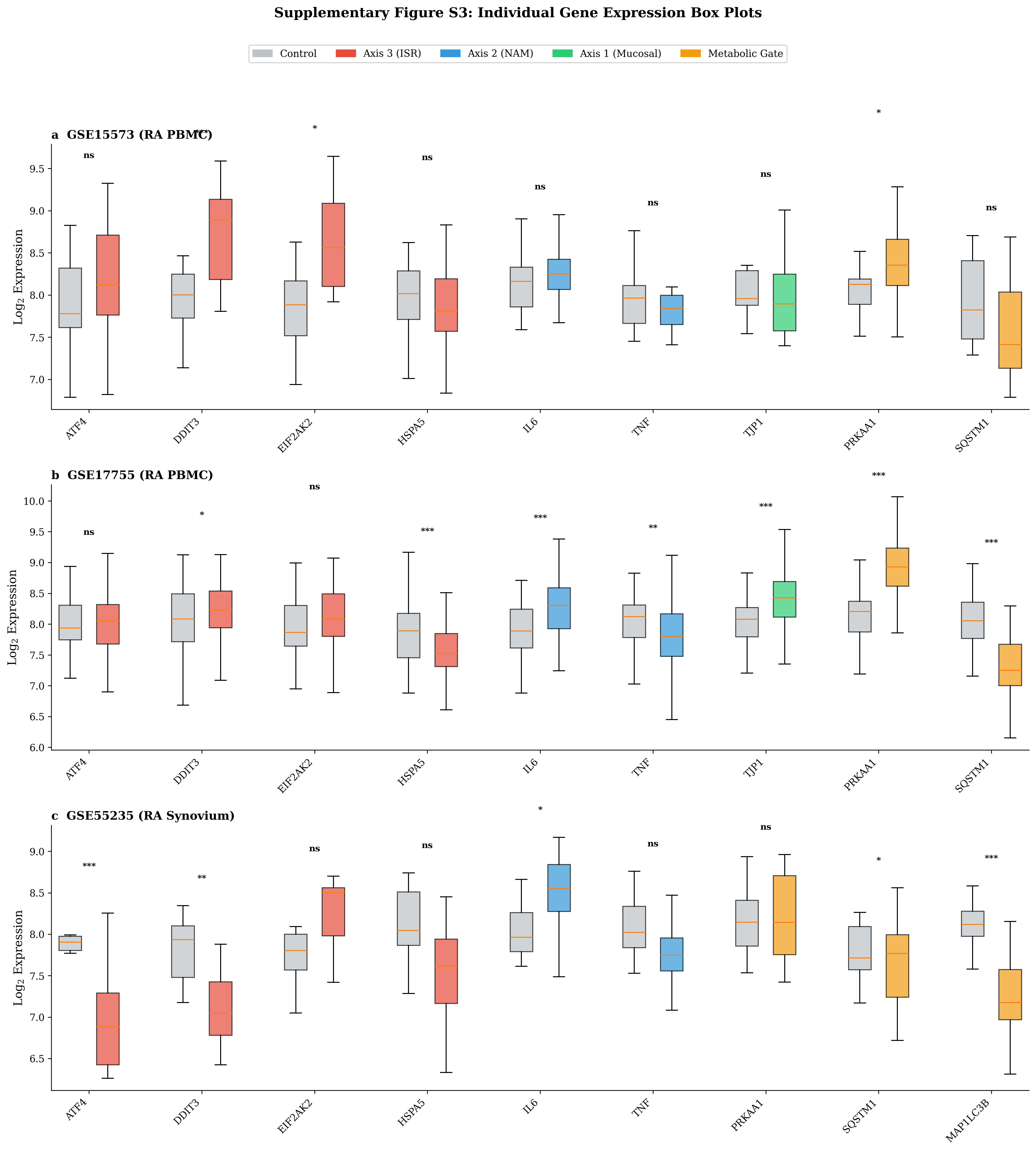

**Supplementary Figure S3.** Individual gene expression box plots. Expression levels (log₂ intensity or RPKM) for all axis-related genes across each dataset, comparing disease versus healthy control groups. Box plots show median, interquartile range, and individual data points. Statistical significance indicated by asterisks (*p<0.05, **p<0.01, ***p<0.001, Benjamini-Hochberg corrected).

### Supplementary Figure S4

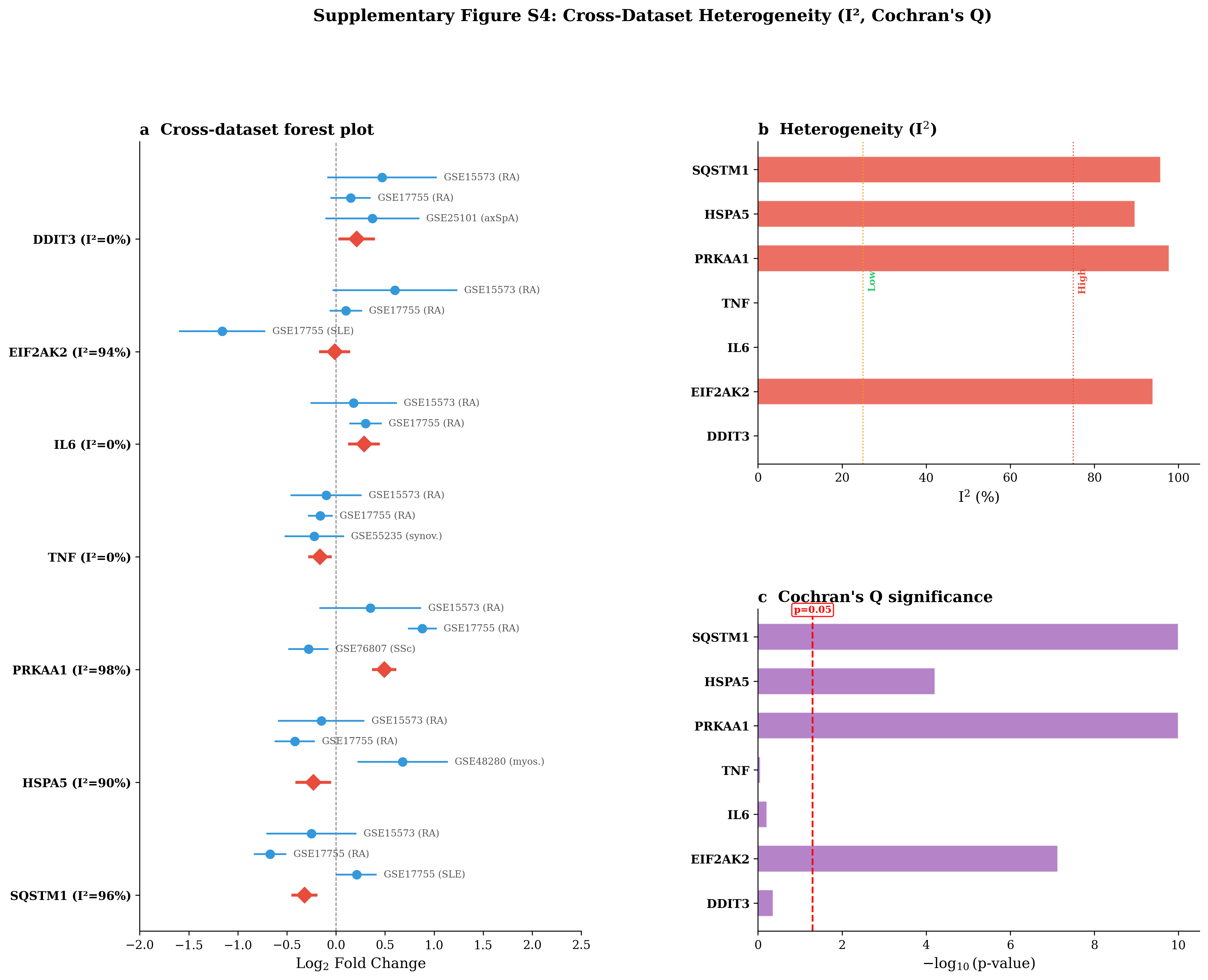

**Supplementary Figure S4.** Extended cross-dataset consistency analysis. (a) Forest plot showing Log₂FC and 95% CI for each gene across all available datasets, with random-effects meta-analytic summary (red diamonds). (b) Heterogeneity assessment using I² statistics with thresholds for low (<25%), moderate (25-75%), and high (>75%) heterogeneity. (c) Cochran's Q test significance (−log₁₀ p-value); dashed line indicates p=0.05 threshold.

### Supplementary Figure S5

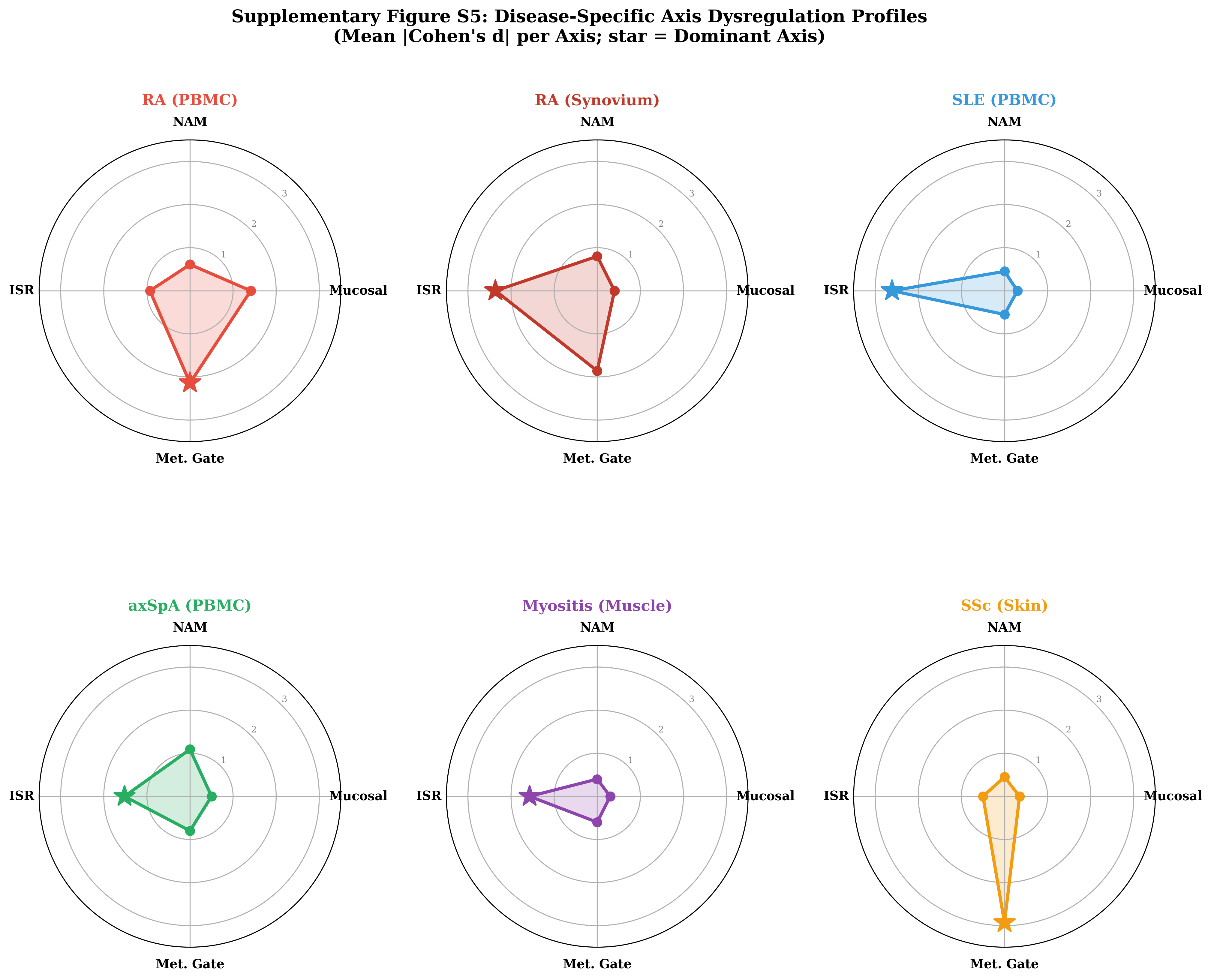

**Supplementary Figure S5.** Disease-specific axis dysregulation radar profiles. Mean absolute Cohen's d per axis for each dataset, displayed as radar plots. Each vertex represents one of the three biological axes plus the metabolic gate. Larger areas indicate greater overall dysregulation. RA blood shows balanced multi-axis involvement, while SLE and myositis show ISR-dominant profiles.
